# Gut microbiota-dependent phenylpropanoic acid derivatives reduced in cancer cachexia protect against myotube atrophy

**DOI:** 10.64898/2026.09.17.752337

**Authors:** Alexandra L. Degraeve, Xiaolin Li, Camille Lefevre, Edwige Piron, Axelle Loriot, Nathalie M. Delzenne, Audrey M. Neyrinck, Laure B. Bindels

**Author notes:** Co-shared first authorship (Alexandra L. Degraeve, Xiaolin Li). **Correspondence:** Laure B. Bindels,. Avenue E. Mounier, 73, B1.73.11, Brussels, Belgium.

## Abstract

Cancer cachexia is a debilitating disease characterized by muscle atrophy. Given the gut dysbiosis in cancer cachexia and the increasing evidence of a gut-muscle axis, we explored the potential beneficial effects of bacteria-dependent metabolites on myotube atrophy. Using both hypothesis-driven and hypothesis-free approaches, in-depth metabolomic analysis of blood samples from cachectic C26 tumor-bearing mice treated or not with antibiotics, as well as disease-free germ-free and conventionalized mice, identified 7 bacteria-dependent metabolites decreased under cachectic conditions. Such alterations were not mediated by reduced caloric intake. Among them, 2 metabolites, namely 2-phenylpropanoic acid (2PPA, also known as 2-phenylpropionic acid) and 3-(3,4-dihydroxyphenyl)propanoic acid (3,4OHPP, also known as 3,4-dihydroxyhydrocinnamic acid or dihydrocaffeic acid), demonstrated anti-atrophying effect, alone and in combination, on mouse C2C12 myotubes. Transcriptomics revealed that these 2 bacteria-dependent metabolites restored the amino acid homeostasis with an activation of ATF4 and the serine biosynthesis pathway. Pharmacological inhibition of the phosphoglycerate dehydrogenase (PHGDH), the rate-limiting enzyme of this pathway, prevented the anti-atrophying effects of 2PPA and 3,4OHPP, indicating a causal role for PHGDH in this effect. By identifying microbiota-dependent metabolites as potential therapeutic levers, the current work not only advances our understanding of microbiome-host crosstalk in disease but also opens avenues for innovative, targeted interventions to mitigate muscle atrophy.

Graphical abstract.Bacteria-dependent metabolites that are decreased upon cancer cachexia in mice mitigate muscle atrophy *in vitro.* 7 metabolites were identified as depleted in cachectic condition and microbiota-dependent, among which 2 of them, namely 2-phenylpropanoic acid (2PPA) and 3-(3,4-dihydroxyphenyl)propanoic acid (3,4OHPP), counteracted C2C12 myotube atrophy induced by dexamethasone. Created with BioRender.com.

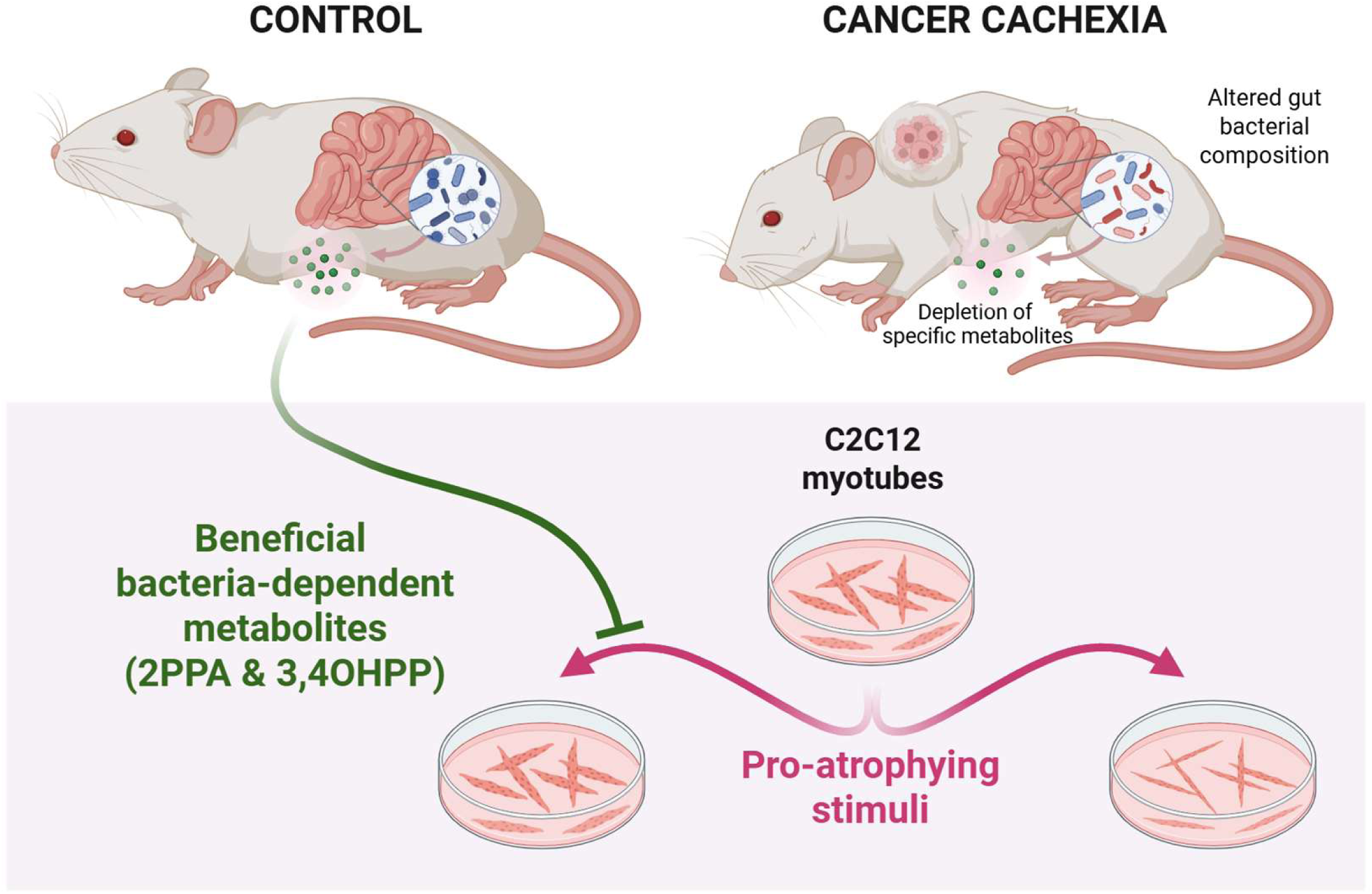

## INTRODUCTION

Cancer cachexia is a multifactorial syndrome characterized by body weight loss, muscle atrophy, fat depletion, anorexia and systemic inflammation^1-3^. Cardiac, bone and liver disturbances are also frequently reported^3,4^. The prevalence of cachexia varies depending on the cancer type, ranging from 15% in prostate cancer up to 45% in colorectal cancer and 65% in pancreatic cancer^5^. Clinically, cancer cachexia results in increased morbidity and mortality rates as well as reduced tolerance to anti-cancer treatments^5^. The best option to treat this multifactorial syndrome appears to be a multimodal approach targeting the multiple components of cachexia (e.g. muscle wasting, fat mass loss, anorexia, fatigue) and consisting in nutritional support, physical exercise, and pharmacological agents^6-9^. However, the nature of the pharmacological and nutritional tools remains a matter of debate^10-12^.

The mechanistic understanding of cancer cachexia is currently incomplete. The prevailing consensus is that several tumor- and host-derived pro-inflammatory mediators and catabolic factors drive the communications between tissues such as the tumor, muscles, adipose tissues, and the liver^3,13-15^. In this context, muscle atrophy results from an imbalance between protein synthesis and degradation, both processes being regulated by multiple factors, pathways and systems (e.g. IGF-1 and TGF-β signalling pathways, pro-inflammatory cytokines, ubiquitin-proteasome and autophagy-lysosomal systems, oxidative stress and mitochondrial function, glucocorticoids signalling)^16-18^. In this context, a potential contribution of the gut microbiota to muscle metabolism is emerging^13,16,19-21^.

The gut microbiota comprises trillions of microbes that reside in the gastrointestinal tract of mammals and is a key regulator of host metabolism and immunity. The appreciation that the gut microbiome influences health has prompted tremendous interest in the mechanisms underlying this crosstalk^22^. Works from our lab and others have contributed to the establishment of the gut microbiota as innovative therapeutic target in the context of cancer cachexia. Indeed, gut dysbiosis has been associated with cancer cachexia in mice^13,20,23,24^ and patients^25-28^. Furthermore, we showed that restoring the lactobacilli levels through the administration of rationally selected lactobacilli counteracted muscle atrophy and decreased systemic inflammation in cachectic mice^29^. We also established that nutritional interventions that target the microbiota (mainly prebiotics) decreased cancer progression, reduced morbidity, and/or increased survival of cachectic mice^30-32^.

The main metabolites studied in the gut-muscle axis are short-chain fatty acids^21^. However, as treating germ-free mice with short-chain fatty acids only partly reversed the skeletal muscle impairments^33^, we postulated that bacterial metabolites other than short-chain fatty acids, namely bacterial amino acid metabolites (bAAms) but also unknown metabolites, may be involved in the gut-muscle axis. This hypothesis of a role for bAAms is supported by our metabolomics study which highlighted an increase in the fecal levels of most amino acids, that we attributed to both an increased transit time and a reduced bacterial transformation^34^. Targeted analyses of the bacterial metabolites of tryptophan further confirmed this hypothesis of a reduced amino acid fermentation^35^. Based on these data, we hypothesized that the reduced bacterial metabolism of AA contributes to the muscle wasting in cancer cachexia, and reciprocally that bAAms can support muscle mass and function.

Therefore, in the current work, we aimed to further determine the contribution and therapeutic interest of bAAms and other similar compounds to muscle wasting. To do so, chemical isotope labelling liquid chromatography–mass spectrometry (CIL-LC-MS) metabolomics was applied to the systemic blood of healthy and cachectic mice with or without microbiota depletion. Such analyses revealed a reduction in 7 bacteria-dependent metabolites, including 2 established bAAms. Among these 7 metabolites, 2 (namely, 2-phenylpropanoic acid [2PPA] and 3-(3,4-dihydroxyphenyl)propanoic acid [3,4OHPP]) attenuated myotube atrophy *in vitro*. Whole transcriptome analysis of myotubes was performed followed by pharmacological targeting to explore the mechanism of action underlying the beneficial effects of these 2 bacteria-dependent metabolites.

## RESULTS

### Seven bacteria-dependent metabolites are reduced in cachectic mice independently of anorexia

As our aim was to systematically investigate the potentially beneficial role of bAAms and small polar compounds other than short-chain fatty acids, the systemic blood of healthy and cachectic mice with or without microbiota depletion was analyzed using MS-based metabolomics following the protocol depicted in Figure 1A. The well-characterized Colon 26 carcinoma (C26) model was used and consists of the ectopic implantation of mouse C26 cells in mice leading to a small reproductible tumor growth inducing cachexia^36-38^. Pair-fed (PF) mice, receiving the amount of food consumed by CT or C26 mice, were included in the analysis to isolate the effect of anorexia.

**Figure 1.**
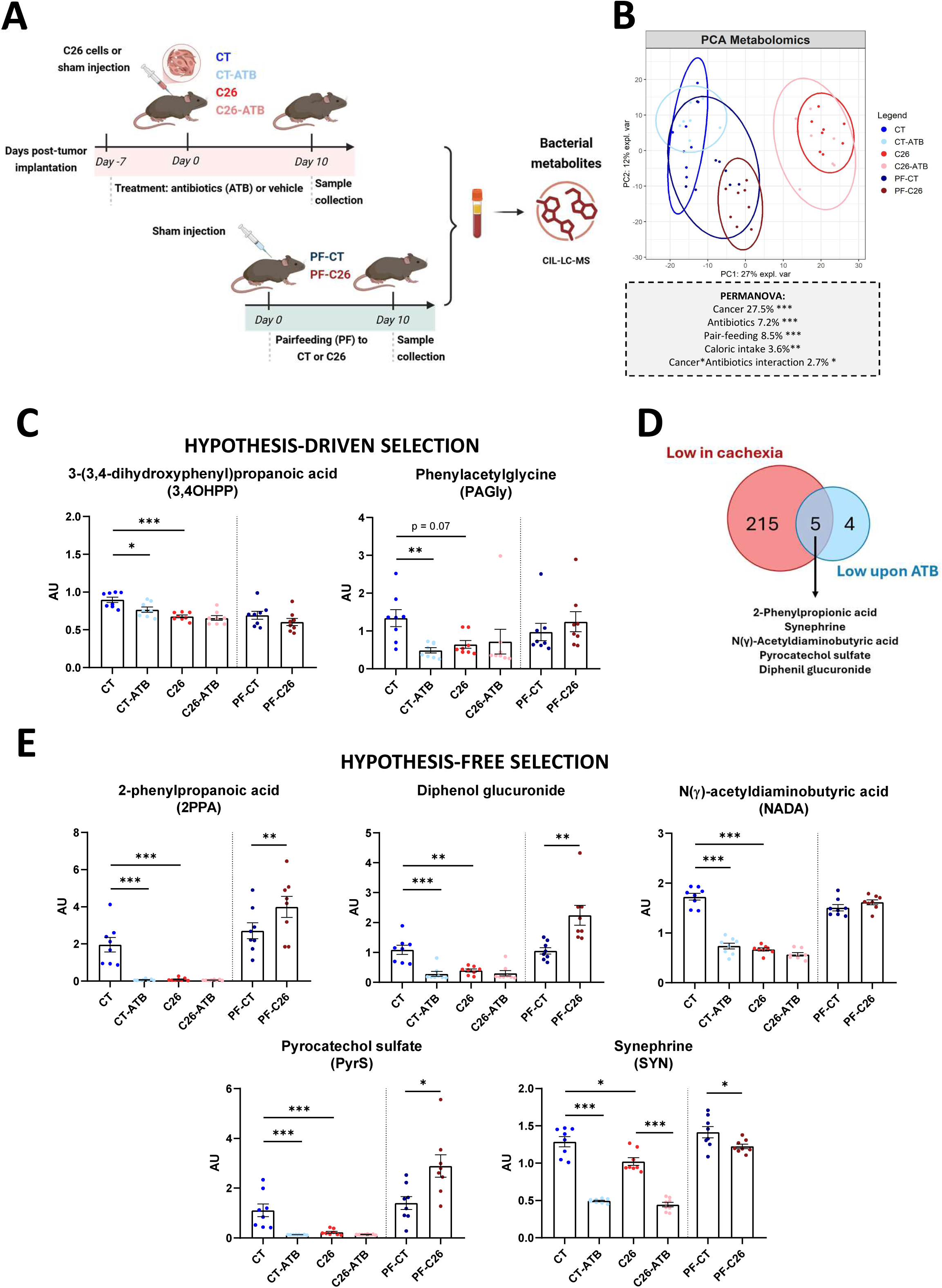
Identification and selection of bacteria-dependent metabolites decreased in tumor-bearing C26 mice. (**A**) Schematic overview of the experimental design. (**B**) Principal component analysis (PCA) of the levels of circulating metabolites across experimental groups. In the PERMANOVA analysis, the pair-feeding effect refers to the fact that some experimental groups (namely PF-CT and PF-C26) were restricted in the amount of food that were given while the caloric intake effect reflects the fact that some experimental groups (C26, C26-ATB and PF-C26) consumed less calorie due to anorexia or per design. (**C**) Relative concentration of metabolites selected based on a hypothesis-driven approach: 3-(3,4-dihydroxyphenyl)propanoic acid (3,4OHPP) and phenylacetylglycine (PAGly). (**D**) Venn diagram illustrating metabolites that were reduced in C26 mice and in ATB mice. Metabolites were classified as “low in cancer” if significantly decreased in C26 mice compared with controls, and as “low upon antibiotics” if significantly decreased in ATB mice compared with vehicle controls (q < 0.05, Tukey’s post hoc test). (**E**) Relative concentration of metabolites selected using a hypothesis-free approach: 2-phenylpropanoic acid (2PPA), diphenol glucuronide, N(γ)-acetyldiaminobutyric acid (NADA), pyrocatechol sulfate (PyrS), and synephrine (SYN). Data represented in **C, E** were analyzed by two-way ANOVA with Tukey’s post hoc test on log-transformed data for evaluating the ATB and C26 effects, while Student t-test on log-transformed data was used for comparing PF-CT vs PF-C26. AU, arbitrary units. N = 8 mice per group. \**p* ≤ 0.05, \*\**p* ≤ 0.01, and \*\*\**p* ≤ 0.001. AU: arbitrary units.

Multivariate analysis revealed a major impact of cancer on the blood metabolome with a minor impact of microbiota depletion by antibiotics (ATB, Figure 1B). Specifically, tumoral status explained 27.5% of the variance in the dataset while antibiotherapy accounted for 7.2% of the variance, with an interaction between both parameters, indicating that the impact of antibiotics was not the same in healthy mice and in cachectic mice. In this model, we also integrated the reduced caloric intake and the pair-feeding effects. The caloric intake effect reflects the fact that some experimental groups (C26, C26-ATB and PF-C26) consumed less calories due to anorexia or per design, while the pair-feeding effect refers to the fact that some experimental groups (namely PF-CT and PF-C26) were restricted in the amount of food that they were given. Reduced caloric intake and pair-feeding had minor impacts on the blood metabolome, respectively explaining 3.6% and 8.5% of the variance in the dataset. We therefore concluded that reduced caloric intake *per se* does not drive the alterations in the metabolome observed in C26 cachectic mice.

Following up with univariate analyses, we applied a hypothesis-driven approach. Namely, we explored the impact of cachexia and antibiotics on the levels of bAAms identified through a literature review^20,39,40^ and chemical analogs were selected too (list of metabolites in Supplemental Table 1). Among the 90 screened metabolites, the levels of only 2 metabolites, namely 3-(3,4-dihydroxyphenyl)propanoic acid (3,4OHPP, also known as 3,4-dihydroxyhydrocinnamic acid or dihydrocaffeic acid) and phenylacetylglycine (PAGly) were reduced both upon antibiotics exposure and in cachectic mice (Figure 1C). These reductions were not explained by the reduced caloric intake. As the levels of these 2 metabolites were not further reduced in cachectic mice upon antibiotics administration, we concluded that their bacterial production was completely blunted in cachectic mice. Next, we carried out a hypothesis-free analysis where all metabolites were considered. After correction for multiple testing, 5 metabolites were identified as reduced in cancer cachexia and upon microbiota depletion, namely 2-phenylpropanoic acid (2PPA, also known as 2-phenylpropionic acid), diphenol glucuronide, N(γ)-acetyldiaminobutyric acid (NADA), pyrocatechol sulfate (PyrS), and synephrine (SYN) (Figure 1D). Caloric restriction partially recapitulates the reduction in SYN, while it was not the case for the other compounds. SYN levels were further reduced in microbiota-depleted cachectic mice compared to cachectic mice with intact microbiota, indicating that the bacterial production of SYN was only partially affected by cachexia. On the opposite, the levels of 2PPA, NADA, PyrS and diphenol glucuronide were not further reduced in cachectic mice upon antibiotics, indicating that their bacterial production was completely blunted in cachectic mice (Figure 1E).

To validate that these 7 compounds are bacteria-dependent metabolites, we investigated the impact of the conventionalization of germ-free (GF) mice with a complex gut microbiota on their blood level following the design depicted in Figure S1A. Consistent with the antibiotic effect, PAGly level was significantly decreased in the absence of gut microbiota (GF) as compared to the conventionalized mice (CVZ), and similar trend was observed for 3,4OHPP (Figure S1B). All metabolites identified by the hypothesis-free approach depicted reduced blood level in GF mice, reaching significance for NADA and SYN (Figure S1C).

Altogether, this analysis led us to the identification of 7 bacteria-dependent metabolites that are reduced in cachectic mice independently of anorexia.

### Two bacteria-dependent metabolites, namely 2PPA and 3,4OHPP, exert anti-atrophic effects in myotubes

Next, to explore the beneficial potential of such compounds on muscle atrophy, we exposed fully differentiated C2C12 murine myotubes to each compound individually (except for diphenol glucuronide, not commercially available) in the presence of dexamethasone (Figure 2A). Dexamethasone is a synthetic glucocorticoid widely used to induce skeletal muscle atrophy *in vitro* and to reproduce several catabolic pathways implicated in cancer cachexia^41^. We assessed the impact of each bacterial compound on myotube diameter at a concentration of 100 µM. None of the compounds induced myotube cytotoxicity (Figure S2). As expected, dexamethasone reduced the myotube diameter (Figure 2B). Among the tested compounds, only 2PPA and 3,4OHPP prevented dexamethasone-induced myotube atrophy.

**Figure 2.**
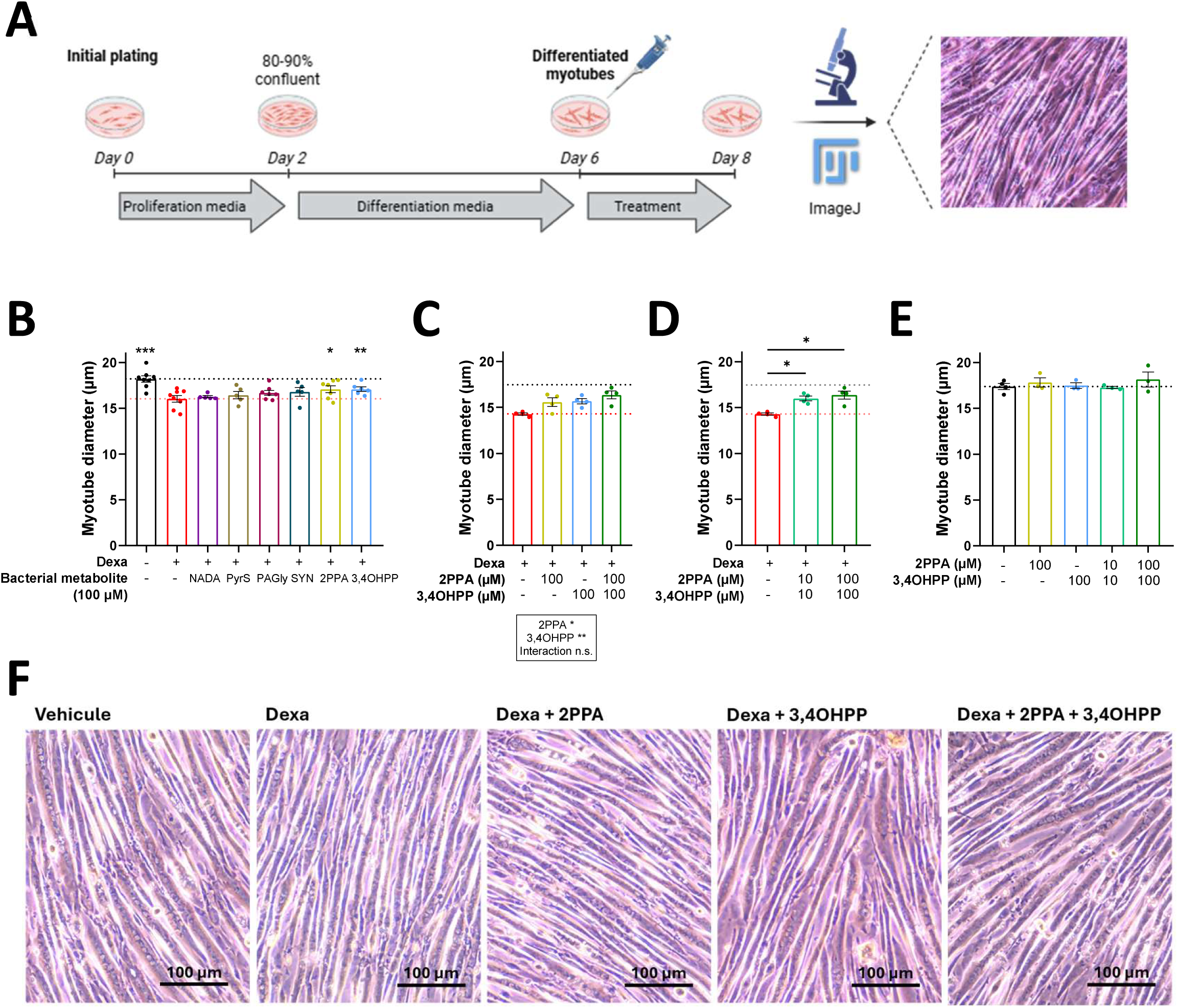
Selected bacteria-dependent metabolites attenuate dexamethasone-induced muscle atrophy in C2C12 myotubes. (**A**) Schematic overview of the *in vitro* C2C12 experimental protocol. After 48 h of treatment, myotubes diameter was measured in each condition. (**B**) Treatment with 100µM of individual candidate metabolites (N(γ)-acetyldiaminobutyric acid [NADA], pyrocatechol sulfate [PyrS], phenylacetylglycine [PAGly], synephrine [SYN], 2-phenylpropanoic acid [2PPA], 3-(3,4-dihydroxyphenyl)propanoic acid [3,4OHPP]) in the presence of 1 µM dexamethasone (Dexa). N = 5-8 experiments, n = 3 technical replicates. (**C**) Treatment with 2PPA and/or 3,4OHPP in the presence of 1 µM Dexa. N = 4 experiments, n = 3 technical replicates. (**D**) Treatment with or without combined 2PPA and 3,4OHPP at 10 or 100 µM, in the presence of 1 µM Dexa. N = 4 experiments, n = 3 technical replicates. **C** and **D** are reporting the same results obtained for 2 conditions (Dexa, 2PPA and 3,4OHPP (100 µM)). (**E**) Treatment with 2PPA and/or 3,4OHPP at 10 or 100 µM, in the absence of Dexa. N = 1 experiment, n= 3 technical replicates. In **B**-**E**, black and red horizontal dotted lines indicate the mean value of the vehicle control and Dexa conditions, respectively, (**F**) Representative pictures of phase contrast microscopy of myotubes treated with the different conditions presented in **C**. Data were analyzed using a paired t-test compared with Dexa alone in **B**, with a two-way ANOVA in **C**, with a one-way ANOVA with Dunnett’s post-test as compared to Dexa in **D** and to vehicle control in **E**. \**p* ≤ 0.05, \*\**p* ≤ 0.01, and \*\*\**p* ≤ 0.001.

Then, simultaneous evaluation of the impact of 2PPA and 3,4OHPP, individually and in combination, revealed an absence of interaction indicating that their anti-atrophic effects are not synergic (aka, the combined effect is not superior to the addition of each individual effect). However, dexamethasone-induced atrophy was more strongly counteracted by the combination than by each compound alone (2PPA + 3,4OHPP corrected the dexamethasone effect by 65.4%, 2PPA by 40.3%, and 3,4OHPP by 43.9%; Figure 2C, F). We thus decided to continue exploring the combination. As the levels of bAAms are often in the range of 10 µM or below^35,40,42^, we also tested the impact of 10 µM of 2PPA and 10 µM of 3,4OHPP on myotube diameter. The combination of each compound at 10 µM was sufficient to prevent myotube atrophy to a similar extent than the combination of each compound at 100 µM (Figure 2D). Finally, incubation of myotubes with 2PPA and 3,4OHPP in the absence of atrophying stimuli did not lead to an increase in myotube diameter, leading us to conclude that, while these compounds have anti-atrophic properties, they do not exhibit protrophic action at these levels (Figure 2E).

### Combination of 2PPA and 3,4OHPP counteracts alterations in gene networks related to amino acid homeostasis under the control of ATF4

To explore the mechanism by which the combination of 2PPA and 3,4OHPP can counteract myotube atrophy, we performed whole transcriptome analysis of the myotubes under the different conditions, namely vehicle, dexamethasone, and combination of 2PPA and 3,4OHPP at 10 µM and 100 µM in the presence of dexamethasone. Such analysis revelated a major impact of dexamethasone on the myotube transcriptome as evidenced from the shift alongside the first component on the principal component analysis (PC1 90%, Figure 3A). The impact of the combination was evidenced alongside the second component (PC2 5%), with a stronger effect at 100 µM than at 10 µM. These findings align with the number of genes found to be affected in each condition. Dexamethasone affected the levels of 7167 genes, with 3662 genes being upregulated and 3505 genes being downregulated (Figure 3B). At 10 µM, the combination of 2PPA and 3,4OHPP altered the expression of 328 genes (191 genes up and 137 genes down), while at 100 µM, the combination altered the expression of 1610 genes (836 genes up and 774 genes down) (Figures 3C and 3D).

**Figure 3.**
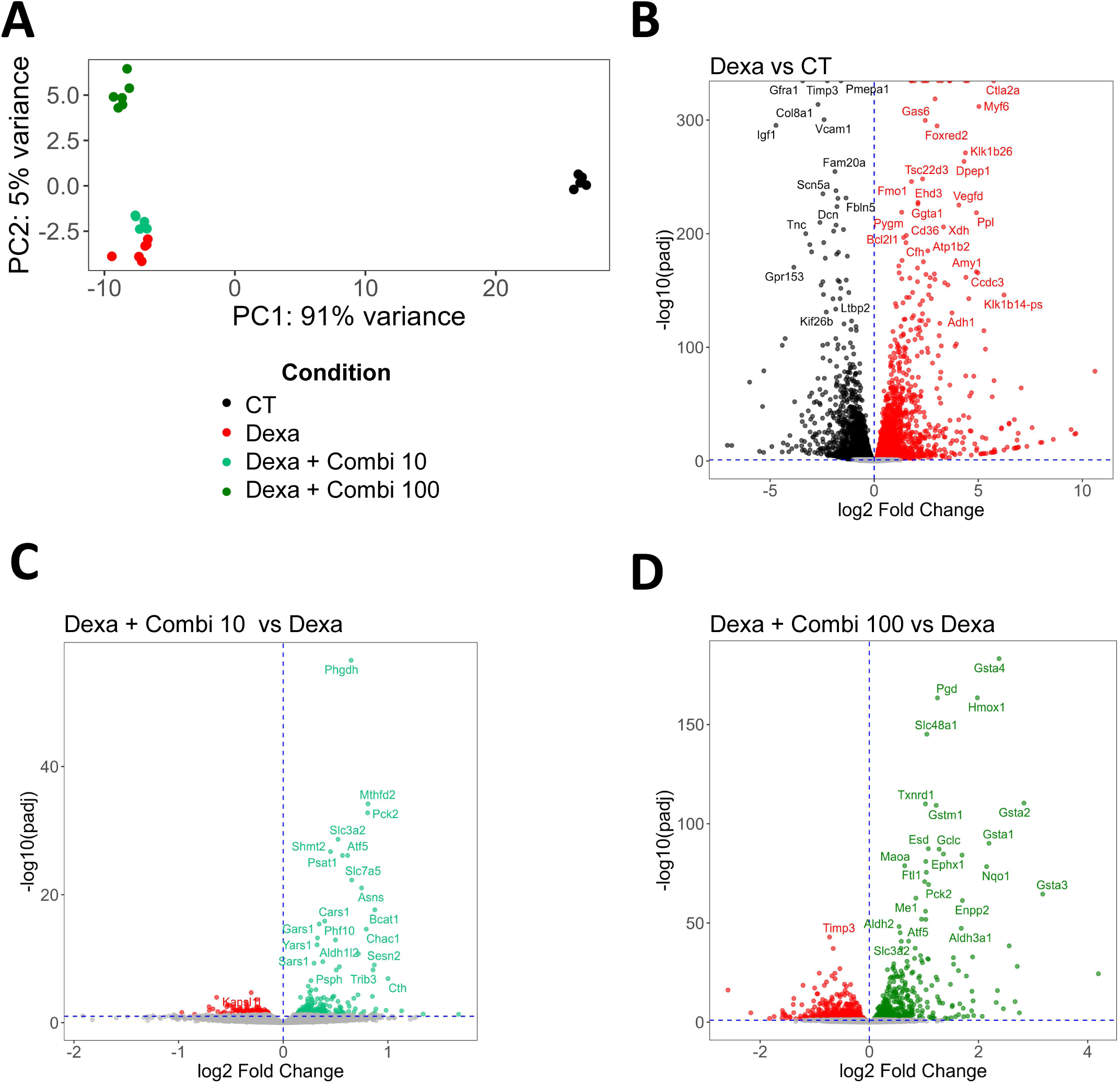
Combination of 2PPA and 3,4OHPP alters myotube transcriptome. (**A**) Principal component analysis (PCA) of gene counts in myotubes following combined treatment with 2PPA and 3,4OHPP at 10 or 100 µM in the presence of 1 µM dexamethasone. (**B**-**D**) Volcano plots showing the results of differential gene expression analyses across different conditions. Grey dots represent non-differentially expressed genes (padj > 0.05), colored dots represent genes significantly upregulated in each condition: under dexamethasone (Dexa) alone (red) versus vehicle (CT, black) condition (**B**), under combined treatment with 2PPA and 3,4OHPP at 10 µM in the presence of Dexa (Combi 10, light green) versus Dexa alone (red) (**C**), under combined treatment with 2PPA and 3,4OHPP at 100 µM in the presence of Dexa (Combi 100, dark green) versus Dexa alone (red) (**D**). Blue horizontal dotted line indicates the significance threshold (padj < 0.05). N = 1 experiment, n= 6 technical replicates.

As the anti-atrophic effect of the combination of 2PPA and 3,4OHPP was similar at the two concentrations, we reasoned that the modifications found for the condition 10 µM would be enough to elicit the phenotype. We therefore focused our analysis on these samples. Over-representation analysis on the 328 genes altered by the combination at 10 µM revealed changes in many pathways related to amino acid homeostasis (transport and metabolism of amino acids, including the synthesis of glycine, serine and threonine), aminoacyl tRNA (tRNA aminoacylation and biosynthesis), metabolism of folate and the ATF4 pathway (REACTOME and KEGG gene sets, Figures 4A and 4B, respectively). A prediction of the transcription factors involved in the regulation of the genes upregulated and downregulated by the combination at 10 µM was carried out. No transcription factors were identified when looking at the 137 downregulated genes. Eleven significant transcription factors were identified when focusing on the 191 upregulated genes, with ATF4 at the top of the ranking (Figure 4C). As ATF4 is a key transcriptional regulator of the genes involved in the serine and glycine synthesis pathway^43^, we further explored the enrichment of the KEGG pathway “Glycine, serine and threonine metabolism”. We found out that 5 genes (i.e., *Phgdh*, *Shmt2*, *Psat1*, *Cth*, *Psph*) involved in the biosynthesis of serine and cysteine were upregulated in this KEGG pathway, including *Phgdh*, coding for the phosphoglycerate dehydrogenase, the rate-limiting enzyme for serine biosynthesis^44,45^ (Figures 4D and 4E).

**Figure 4.**
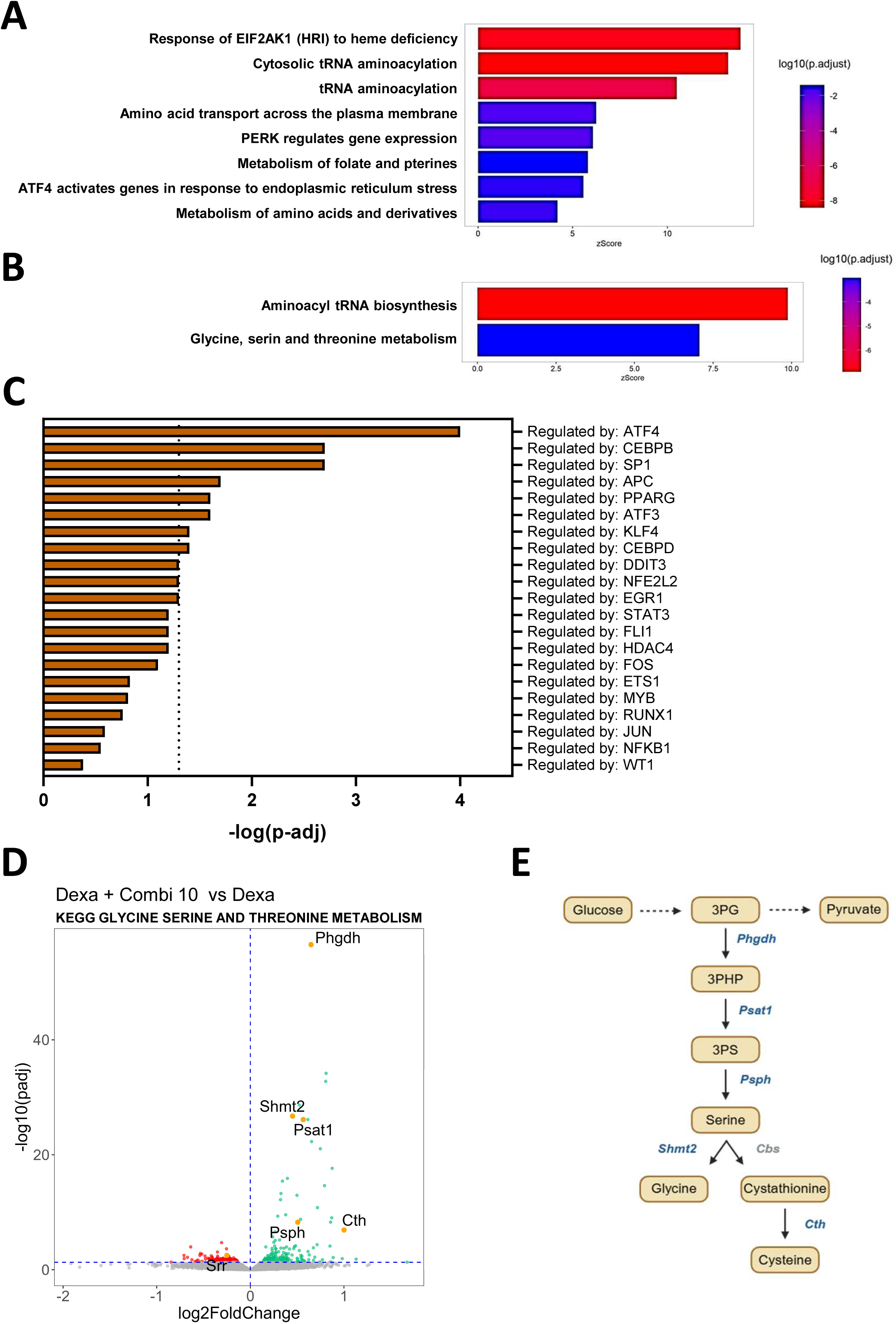
Combination of 2PPA and 3,4OHPP activates metabolic pathways related to amino acid homeostasis. (**A**) Significantly enriched REACTOME pathways between myotubes treated with 2PPA and 3,4OHPP at 10 µM in the presence of 1 µM Dexa compared to Dexa-treated myotubes identified through over-representation analysis (ORA). (**B**) Significantly enriched KEGG pathways between myotubes treated with 2PPA and 3,4OHPP at 10 µM in the presence of 1 µM Dexa and Dexa-treated myotubes identified through ORA. (**C**) Transcription factors predicted to be involved in the regulation of the 191 genes significantly upregulated in myotubes treated with 2PPA and 3,4OHPP at 10 µM in the presence of 1 µM Dexa compared to Dexa-treated myotubes using TTRUST implemented in Metascape. All transcription factors with a p-value < 0.05 are depicted. The vertical dotted line indicates the significant threshold (p-adj ≤ 0.05). (**D**) Volcano plot showing the results of differential gene expression analysis when comparing combined treatment with 2PPA and 3,4OHPP at 10 µM in the presence of Dexa (Combi_10, light green) versus Dexa alone (red). Grey dots represent non-differentially expressed genes (padj > 0.05), colored dots represent significantly differentially expressed genes (padj < 0.05, blue horizontal dotted line). Orange dots are genes from the KEGG pathway “Glycine, serine and threonine metabolism” that are differentially expressed (padj < 0.05). (**E**) Diagram depicting the metabolic function of the genes upregulated by the treatment with 2PPA and 3,4OHPP at 10 µM (in blue) within the KEGG pathway “Glycine, serine and threonine metabolism”. N = 1 experiment, n= 6 technical replicates. 3PG: 3-phosphoglycerate; 3PHP: 3-phosphohydroxypyruvate; 3PS: 3-phosphoserine; *Phgdh*: phosphoglycerate dehydrogenase; *Psat1*: phosphoserine aminotransferase 1; *Psph*: phosphoserine phosphatase; *Shmt2*: serine hydroxymethyltransferase 2; *Cbs*: cystathionine β-synthase; *Cth*: cystathionine γ-lyase.

To test the robustness of these results, we identified in a separate analysis the genes affected by dexamethasone and modified in the opposite way by both concentrations of the combination of 2PPA and 3,4OHPP. This approach allowed us to pinpoint 77 genes (18 downregulated and 59 upregulated) (Figure S3A). Over-representation analysis on these 77 genes revealed similar changes, namely changes in pathways related to amino acid homeostasis, to aminoacyl tRNA and to the GCN2/ATF4 pathways (REACTOME and KEGG gene sets, Figures S3B and S3C, respectively). No transcription factors were identified when looking at the 18 downregulated genes. Six transcription factors were identified when considering the 59 upregulated genes, with again ATF4 at the top of the ranking (Figure S3D).

Altogether, these data indicate that the combination of 2PPA and 3,4OHPP prevent alterations in gene networks related to amino acid homeostasis and tRNA acylation, and pinpoint ATF4 as a key transcription factor involved in this transcriptome regulation.

### The anti-atrophic effect of the combination of 2PPA and 3,4OHPP may be mediated by PHGDH

As ATF4 regulates the expression of the genes involved in the control of serine and glycine synthesis, a pathway that has been shown to be essential in basal conditions for myotube growth^46-48^, we hypothesized that the combination of bacteria-dependent metabolites may exert its beneficial impact through the activation of ATF4 and the stimulation of the downstream serine biosynthesis pathway. To test this hypothesis, myotubes were incubated in presence or absence of the PHGDH inhibitors CBR-5884^49^ and NCT-503^50^. A compound structurally related to NCT-503 but pharmacologically inactive against PHGDH was included, namely ‘PHGDH-inactive’.

As expected, the reduction of myotube diameter by dexamethasone was counteracted in the presence of combined 2PPA and 3,4OHPP at 100 µM. The anti-atrophic effect of the combination of bacteria-dependent metabolites was lost upon incubation with CBR-5884 at 2.5 and 10 µM (Figure 5A). Then, evaluation of the impact of CBR-5884 itself on normal or atrophic myotubes indicated that the inhibitor has no intrinsic activity affecting myotube diameter (Figure 5B). None of the conditions induced myotube cytotoxicity, except CBR-5884 at 33 µM, a concentration corresponding to the IC50 that was initially included in the evaluation and thus not further used during myotube diameter analysis (Figure S4A-B).

**Figure 5.**
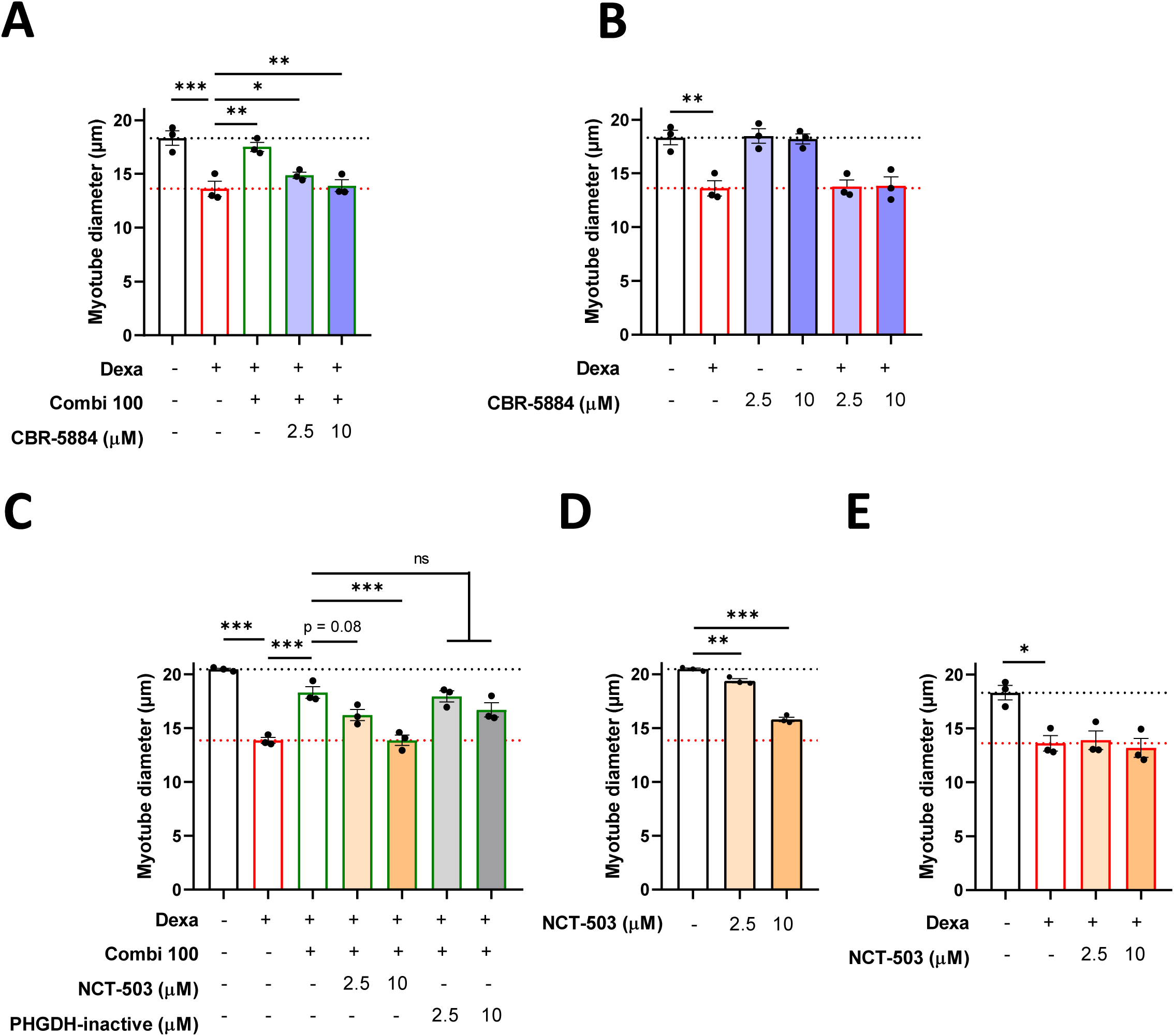
Inhibition of PHGDH attenuates the protective effect of combined metabolites against dexamethasone-induced myotube atrophy. C2C12 myotubes were treated with different conditions for 48 h and myotubes diameter was measured. (**A**) In the presence or absence of 1 µM dexamethasone (Dexa) combined or not with 2PPA and 3,4OHPP (Combi 100 µM) and the PHGDH inhibitor CBR-5884 (2.5, 10 µM). N = 3 experiments, n = 3 technical replicates. (**B**) In the presence or absence of 1 µM Dexa, and with or without the PHGDH inhibitor CBR-5884 (2.5, 10 µM). N = 3 experiments, n = 3 technical replicates. **A** and **B** are reporting the same results obtained for 2 conditions (Vehicle, Dexa). (**C**) In the presence or absence of 1 µM Dexa, combined or not with Combi 100 µM, with or without the PHGDH inhibitor NCT-503 (2.5, 10 µM), and with or without PHGDH-inactive (2.5, 10 µM). PHGDH-inactive is a compound structurally related to NCT-503, but pharmacologically inactive against PHGDH. N = 3 experiments, n = 3 technical replicates. (**D**) In the presence or absence of the PHGDH inhibitor NCT-503 (2.5, 10 µM). N = 3 experiments, n = 3 technical replicates. **C** and **D** are reporting the same results obtained for Vehicle. (**E**) In the presence or absence of 1 µM Dexa, combined or not the PHGDH inhibitor NCT-503 (2.5, 10 µM). N = 3 experiments, n = 3 technical replicates. Black dotted line represents the mean value of the vehicle control condition, red dotted line represents the mean value of the Dexa condition. Data were analyzed using one-way ANOVA with Sidak’s multiple comparisons test. \**p* ≤ 0.05, \*\**p* ≤ 0.01, and \*\*\**p* ≤ 0.001.

In line with the observations obtained with CBR-5884, the combination of metabolites failed to counteract the pro-atrophic effect of dexamethasone on myotubes when co-incubated with NCT-503, another PHGDH inhibitor. In contrast, the ‘PHGDH-inactive’ compound did not interfere with the anti-atrophic action of 2PPA and 3,4OHPP (Figure 5C). When tested alone, NCT-503 induced a dose-dependent reduction in myotube diameter; however, this effect was less pronounced than that of dexamethasone (Figure 5D) and was absent when combined with dexamethasone (Figure 5E). None of these conditions induced myotube cytotoxicity (Figure S4C-E).

Altogether, these results support a role for PHGDH in mediating the anti-atrophic effect of the combination of 2PPA and 3,4OHPP.

## DISCUSSION

Bacterial metabolites are key players in the regulation of host metabolism and immunity by the gut microbiota, with short-chain fatty acids and bile acids being the most studied bacterial (co)metabolites so far. A few bAAms have been identified as mediators of the microbiota-host crosstalk, such as indoles^51-53^, imidazole propionate^54^, phenylacetate^55^ and phenylacetylglutamine^40^. Compounds arising from the bacterial transformation of other types of nutritional compounds are also emerging, such as urolithin A, ensuing from the bacterial transformation of polyphenols^56^ and able to improve mitochondrial function and muscle mass^57^. In this work, we used both a hypothesis-driven approach and a hypothesis-free approach to identify bacteria-dependent metabolites, including bAAms, reduced in cancer cachexia. Among the 7 metabolites identified, 2 of them are known to be bAAms, namely PAGly (derived from phenylalanine^40^) and NADA (derived from aspartate^58^).

Although a contribution of the gut microbiota to the systemic levels of these metabolites is supported by their increased levels in conventionalized mice and their reduction upon antibiotics administration, it remains unclear, at least for some compounds, whether they are bacterial products or ensuing from the impact of the gut microbiome on host enzymes.

Three bacterial pathways for the production of 3,4OHPP have been described, firstly through the transformation of phenylalanine (reported in *Rhodobacter sphaeroides*, an environmental bacterium^59^) suggesting that 3,4OHPP can also be considered as a bAAm, secondly through the transformation of caffeic acid (reported in intestinal bacteria such as *Escherichia coli*, *Bifidobacterium lactis*, and *Lactobacillus gasseri*^60^), and thirdly through the metabolism of eriocitrin (a flavonoid from lemon fruit, reported in gut bacteria *Bacteroides distasonis*, *Bacteroides uniformis*, and *Clostridium butyricum*^61^). By which of these pathways 3,4OHPP is produced in the gut remains to be formally established, although a production derived from phenylalanine is the most likely based on the diet provided to the mice.

Production of 2PPA by bacterial or mammalian pathways has been poorly described, with *in vitro* data indicating that 2PPA could be produced from α-methylstyrene by human hepatic tissue^62^. 2PPA is often confused in the literature and in databases with 3-phenylpropanoic acid (3PPA), a bAAm arising from the bacterial metabolism of phenylalanine and tyrosine^63,64^. 2PPA differs from 3PPA in the position of the phenyl group, making it an unlikely metabolite of tyrosine and phenylalanine. Of note, the glycine derivative of 3PPA was detected in our metabolomics analyses and was not affected by cachexia.

While 2PPA levels are drastically reduced in healthy mice receiving antibiotics, suggesting a critical role of the gut microbiome in the production of this metabolite, the reduction of 3,4OHPP levels in healthy mice upon antibiotics is more modest, indicating that the systemic levels of 3,4OHPP are only partially microbiota-dependent. For all compounds, apart from SYN, antibiotics exposure did not reduce further their systemic levels in cachectic mice, suggesting that their bacterial production is largely blunted in cancer cachexia. Pair-feeding controls were included in the metabolomics study (which are often lacking in this type of study), allowing us to conclude that the reduced production in cachectic mice cannot be attributed to the reduced caloric intake induced by anorexia, at the exception of SYN. Although we cannot formally exclude that the reduction in the levels of the bacteria-dependent metabolites identified in this study is the result of the mere presence of the tumor rather than cachexia itself, we have elements arguing against this hypothesis. First, using NC26 mice which serves as tumor-bearing controls to C26 mice, carrying a tumor similar to C26 but lacking cachexia-inducing properties, we evidenced that only 3% of the ASV whose relative abundance levels are changed in the gut microbiota of C26 mice is attributable to the tumoral presence (manuscript under preparation). Second, in a 2026 multi-compartmental metabolomics study using the C26 model, and including NC26 mice, most plasma-associated metabolomics changes were ascribed to cachexia (C26 vs control: 38.2% of altered metabolites), with only 1.7% of metabolites identified as different in their systemic levels between NC26 mice and control mice^15^. In this dataset, 2PPA and 3,4OHPP were not identified, likely due to a lower depth in metabolomics (233 plasma metabolites identified in this study versus 860 in the current study).

The biological function and impact of 2PPA and 3,4OHPP are poorly defined. 2PPA induces peroxisomal proliferation in rats^65^. 3,4OHPP, also named dihydrocaffeic acid, is recognized for its antioxidant and anti-inflammatory properties^66^. Interestingly, 3,4OHPP can be produced by intestinal bacteria from eriocitrin, a flavonoid present in lemon fruit^61^, that showed anti-atrophying effect on mouse muscle^67^. Whether the effect is mediated by the microbiota and rely on the bacterial metabolism of eriocitrin into 3,4OHPP was not investigated in that study.

Transcriptomics analysis of the myotubes treated jointly with 2PPA and 3,4OHPP revealed changes in many pathways related to amino acid transport and metabolism and tRNA aminoacylation. Analysis of transcriptional regulatory networks identified ATF4 as the transcription factor most strongly activated by the combination of metabolites. ATF4 activation could ensue from a deprivation in essential amino acids (EAA), through the GCN2/eiF2α pathway^68^. Indeed, the increased expression of genes involved in tRNA aminoacylation (such as *Aars1, Cars1, Gars1* and *Sars1*) may reflect accumulation of uncharged tRNA resulting from EAA starvation. Such accumulation is a key activator of GCN2^68-70^. Once activated, GCN2 activates eiF2α through phosphorylation, which in turn activates ATF4 translation^71^. In line with a deprivation in EAA, genes involved in amino acid biosynthesis (eg. *Asns*) and import (e.g. *Lat1*, *Scl7a5*), among which many are target genes of ATF4, were increased by the combination of metabolites. In view of these results, we speculate that 2PPA and/or 3,4OHPP may compete with EAA for cellular entrance, leading to intracellular EAA deprivation and activation of the ATF4 pathway.

Looking at the volcano plot in Figure 4D, we noticed that many of the top genes induced at 10µM of 2PPA and 3,4OHPP belongs to the KEGG pathway “Glycine, serine and threonine metabolism”, and more specifically to the serine biosynthesis pathway (namely, *Phgdh, Shmt2, Psat1, Cth* and *Psph*). Interestingly, ATF4 is an activator of the transcription of many of these genes, including *Phgdh*^72,73^. As PHGDH, the rate-limiting step of the serine biosynthesis pathways, is essential for mouse C2C12 and human primary myotube size^47,48^, we hypothesized that 2PPA and 3,4OHPP may mediate their anti-atrophic effect through an ATF4/PHGDH pathway. The rational to select PHGDH rather than ATF4 as a target for further mechanistic investigation is supported by two elements. First, ATF4 is a key regulator of many intracellular processes^71^. Its inhibition is therefore likely to induce many consequences on multiple pathways. By targeting one of the pathways downstream of ATF4, we are narrowing down our mechanistic investigation. Second, ATF4 can exert both proanabolic and procatabolic effects in muscle, likely due to the nature of its heterodimer partner and the duration of ATF4 activation^16,71,74^, potentially complicating the interpretation of the results of an inhibition of ATF4 in presence of 2PPA and 3,4OHPP.

Inhibiting PHGDH using two structurally different pharmacological inhibitors prevented the anti-atrophic effect of 2PPA and 3,4OHPP, suggesting a key role for PHGDH in the beneficial effect of these bacteria-mediated compounds. Difference in the effect of these two inhibitors on myotube diameter was observed in absence of dexamethasone. A plausible explanation for this difference is the higher potency of NCT-503 toward PHGDH compared to CBR-5884 (IC₅₀ = 2.5 ± 0.6 µM and 33 ± 12 µM, respectively)^49,50^. Of note, a concentration of 33µM CBR-5884 was initially included in our experimental design and excluded due to cytotoxicity.

The mechanisms by which PHGDH contributes to myotube size and biomass has been previously explored^47,48^. PHGDH is essential for the biosynthesis of serine, which is both a proteinogenic amino acid and the source of one-carbon units essential for *de novo* purine and deoxythymidine synthesis^50^. In myotubes, Mantyselka et al. reported findings pointing at functions of PHGDH beyond serine synthesis such as mTORC1 regulation and maintenance of the homeostasis in oxidative stress and redox reactions^48^. In cancer cells, Pacold et al. showed that PHGDH inhibition reduced the incorporation into nucleotides of one-carbon units from glucose-derived and exogenous serine. They suggested that one-carbon unit wasting thus may contribute to the efficacy of PHGDH inhibitors^50^.

A role for one-carbon metabolism in muscle atrophy in cancer cachexia is emerging. One-carbon metabolism is primarily organized around two interconnected cyclic pathways: the folate cycle and the methionine cycle^75^. Morigny et al. showed that over-activation of the methionine cycle drives atrophy and hypermetabolism in myotubes^15^, while Lin et al. reported a disruption of the methionine cycle whose alleviation was able to mitigate cancer-associated muscle atrophy^74^. Upon 2PPA and 3,4OHPP supplementation, *Shmt2* and *Mthfd2,* two key enzymes involved in the folate cycle, are increased (Figure 3C), raising the possibility that an increase in folate cycle may contribute to the anti-atrophic effect of these bacteria-dependent metabolites.

Whether PHGDH contributes to muscle mass maintenance, metabolism or regeneration *in vivo* has not been explored to our knowledge. As our work alongside the one of others^47,48^ indicate a beneficial role for PHGDH in muscle models, it raises the concern of potential muscle wasting side effects for its inhibitors. This concern is timely as PHGDH has emerged as a promising therapeutic target in oncology and PHGDH inhibitors are being actively developed for clinical translation^75^. Studies using pharmacological PHGDH inhibitors such as NCT-503 and CBR-5884 were not oriented towards muscle phenotyping. They reported antitumor activity in mouse models, with some of them indicating no treatment-related body-weight loss and overt systemic toxicity^76^. Consistent with this concern, serine and glycine dietary deprivation strengthened cancer-associated weight loss and muscle atrophy in the HT29 and C26 mouse models^77^. Reciprocally, our results open the possibility of a potential microbiome-based mechanism of resistance in cancer cells towards PHGDH inhibitors, which would be particularly relevant in colorectal cancer.

Altogether, our results identified novel bacteria-dependent metabolites decreased in cancer cachexia and that may be involved in the gut-muscle axis through PHGDH. These metabolites likely arise from the bacterial transformation of amino acids and other nutrients. Further studies will be needed to determine their bacterial production pathways and evaluate their therapeutic potential *in vivo.* This could be achieved either through their direct supplementation or the administration of their precursors alongside their producing bacteria.

## MATERIALS AND METHODS

### Mouse experiments

For the cachectic protocol, forty-eight CD2F1 male mice (7 weeks old, Charles River Laboratories, Italy) were kept in specific pathogen-free (SPF) conditions and housed 2 mice per cage in individually ventilated cages with a 12 h light/dark cycle and fed an irradiated chow diet (AO4-10, SAFE, Tecnilab-BMI, The Netherlands). The experiment was composed of 6 groups of 8 mice, that were randomly assigned in each group based on their body weight. The model used to study cancer cachexia is the well-established C26 model, characterized by body weight and fat mass loss as well as muscle atrophy^37,78^. Seven days after ATB (or water) start, either a saline solution (group CT and group CT-ATB) or C26 cells (1 × 10^6^ cells in 0.1 mL saline; group C26 and group C26-ATB) were subcutaneously injected in the upper left flank.

After one-week acclimatization, in adequate treatment groups, ATB were introduced. The ATB cocktail was composed of neomycin (1 g/L), vancomycin (0.5 g/L), and meropenem (0.25 g/L)^79^. It was administered via drinking water and renewed every other day for the entire experiment. The efficacy of the antibiotic cocktail to deplete the gut microbiota was validated by quantification of total bacteria using qPCR as previously described^79^. In the present study, total bacteria were reduced by 3.57-log upon antibiotics exposure.

To evaluate the impact of reduced caloric intake, pair-fed animals were used. These animals received each day the amount of food consumed that day (day post-injection) by their respective group, either the CT group (PF-CT) or the C26 group (PF-C26). Pair-fed mice were subcutaneously injected in the upper left flank with a saline solution on their day 0.

Ten days after injection (corresponding to a cachectic stage for the C26 injected mice)^78,80^, mice were fasted from 7 A.M. to 1 P.M., and blood samples were harvested following anesthesia (isoflurane gas, Abbot, Belgium). Blood was centrifuge at 13,000 g for 3 min at 4 °C and plasma was collected and frozen in liquid nitrogen. All samples were stored at −80 °C until further analyses.

For the conventionalization protocol, male C57Bl6 germ-free (GF) mice were born and raised at the Ghent Germ-free and Gnotobiotic mouse facility (Ghent University, *LA2400451*). They were maintained in a sterile environment under controlled conditions (10-h light–dark cycle) with ad libitum access to water and diet (2018S, Envigo, USA). GF mice were housed and bred in ‘open’ cages in positive-pressure GF isolators. Before colonization, GF mice were exported from isolators to positive-pressure isocages and left to acclimatize for several days.

Conventionalization of GF mice was performed by diluting mouse cecal and colonic contents (∼0.3 g) in 3 mL reduced phosphate-buffered saline with sterile glass microbeads. Intestinal samples were processed in an anaerobic chamber. Tubes were homogenized for 3 min at 30 Hz and then centrifuged at 800 rpm for 1 min to pellet large insoluble material. 0.2 ml of the supernatant was administered by gavage to each GF mouse of the conventionalized (CVZ) group, with 20G disposable plastic feeding tubes. The mouse intestinal samples were obtained from 3 conventionally raised C57Bl6 SPF mice (7 weeks old, Janvier laboratories). The intestinal samples were obtained shortly before colonization and immediately (within 5 min) diluted, then introduced into the GF mice by gavage within 2 h after dilution, under sterile conditions. The remaining solution was frozen at -80°C. The following day, a second gavage was performed to strengthen the colonization. GF mice were colonized at 8-10 weeks old and maintained for 17 days after conventionalization. At the end of the conventionalization period, GF and CVZ mice were euthanized and samples were collected for further analyses.

All the experiments were approved by and performed in accordance with the guidelines of the local ethics committee from the UCLouvain and Ghent University Faculty Medicine and Health Sciences, Belgium. Housing conditions were as specified by the Belgian Law of 29 May 2013, regarding the protection of laboratory animals.

### Metabolomics

Serum from systemic circulation was analyzed using a 4-channel analysis and chemical isotope labelling liquid chromatography–mass spectrometry (CIL LC–MS) as previously described ^81^. CIL LC-MS is a powerful technique for in-depth metabolome analysis with high quantification accuracy. Unlike conventional LC-MS, it analyses chemical-group-based submetabolomes and uses the combined results to represent the whole metabolome. Using differential isotope labelling (e.g., ^12^C-reagent labelled individual samples spiked with a ^13^C-reagent labelled reference or pooled sample, followed by LC-MS analysis of the resultant mixtures), accurate relative quantification of all labelled metabolites in comparative samples can be performed.

#### Sample preparation

Samples were randomized before any procedures to eliminate potential technical variations from sample preparation and instrument drift. The randomized samples were used for the following preparations and analyses. The samples were split into six aliquots for different labelling methods, and preparation of pooled sample. The aliquot for preparing pooled sample from each individual sample (excluding those with minimal volumes) was combined and mixed thoroughly to generate the pooled sample, which was used as the reference. Each individual plasma sample was spun down. Then, 45 μL of LC-MS grade methanol was added to perform protein precipitation. The methanol extract was completely dried after incubation at -20 °C for 30 minutes, then temporarily stored in -80 °C freezer until labelling.

Chemical Isotope Labeling was performed per channel using a Dansyl-labeling kit or a DmPA-labeling kit. For the one aliquot for amine-/phenol-labelling, 12.5 μL of LC-MS grade water was added prior to labelling. The labelling protocol strictly followed the SOP provided in the kit. Briefly, 6.25 μL of buffer reagent (Reagent A) and 18.75 μL of ^12^C_2_-labeling (for the individual samples and the pooled sample) or ^13^C_2_-labeling (for the pooled sample) reagent (Reagent B) was added into samples. The samples were then vortexed, followed by spinning down. The mixtures were incubated at 40 °C for 45 minutes. After that, 3.75 μL of quenching reagent (Reagent C) was added to quench the excessive labelling reagent. The mixtures were incubated at 40 °C for another 10 minutes. Finally, 15 μL pH adjusting reagent (Reagent D) was added. For the one aliquot of sample for carboxyl-labelling, 12.5 μL of LC-MS grade ACN/water (3:1 v/v) was added prior to labelling. The labelling protocol strictly followed the SOP provided in the kit. Briefly, 5 μL of catalysing reagent (Reagent A) and 25 μL of ^12^C_2_-labeling (for the individual samples and the pooled sample) or ^13^C_2_-labeling (for the pooled sample) reagent (Reagent B) was added into samples. The samples were then vortexed, followed by spinning down. The mixtures were incubated at 80 °C for 60 minutes. After that, 20 μL of quenching reagent (Reagent C) was added to quench the excessive labelling reagent. The mixtures were incubated at 80 °C for another 30 minutes and the chemical isotope labelling procedure was complete. For the one aliquot of sample for hydroxyl-labelling, 12.5 μL of LC-MS grade ACN/water (3:1 v/v) was added prior to labelling. The labelling protocol strictly followed the SOP provided in the kit. Briefly, 12.5 μL of reaction-activating reagent (Reagent A) and 20 μL of ^12^C_2_-labeling (for the individual samples and the pooled sample) or ^13^C_2_-labeling (for the pooled sample) reagent (Reagent B) was added into samples. The samples were then vortexed followed by spinning down. The mixtures were incubated at 60 °C for 60 minutes. After that, 2.5 μL of quenching reagent (Reagent C) was added to quench the excessive labelling reagent. The mixtures were incubated at 60 °C for another 10 minutes. Finally, 12.5 μL of pH adjusting reagent (Reagent D) was added. For the one aliquot of sample for carbonyl-labelling, 12.5 μL of LC-MS grade water was added prior to labelling. The labelling protocol strictly followed the SOP provided in the kit. Briefly, 12.5 μL of pH adjusting reagent (Reagent A) and 12.5 μL ^12^C_2_-labeling (for the individual samples and the pooled sample) or ^13^C_2_-labeling (for the pooled sample) reagent (Reagent B) was added into samples. The samples were then vortexed, followed by spinning down. The mixtures were incubated at 40 °C for 60 minutes, followed by placing in a -80 °C freezer for 10 minutes to stop the reaction. After that, the solution was dried by placing under a nitrogen blower. Finally, the metabolites were re-suspended in 50 μL of LC-MS grade ACN/water (50:50 v/v).

The ^12^C_2_-labeled individual sample was mixed with ^13^C_2_-labeled reference sample in equal volume. The mixture was ready to be analyzed by LC-MS. Prior to LC-MS analysis of the entire sample set, quality control (QC) sample was prepared by equal volume mix of a ^12^C-labeled and a ^13^C-labeled pooled sample.

### LC-MS Analysis Conditions

LC-MS analysis was performed using a Thermo Scientific Vanquish LC system coupled to a Bruker Impact II quadrupole time-of-flight (QTOF) mass spectrometer. Chromatographic separation was achieved on an Agilent Eclipse Plus reversed-phase C18 column (150 × 2.1 mm, 1.8 μm particle size). The mobile phase consisted of (A) 0.1% (v/v) formic acid in water and (B) 0.1% (v/v) formic acid in acetonitrile The elution was carried out using a linear gradient as follows: 25% B at 0 minute, increased to 99% B at 10 minutes, held at 99% B until 15 minutes, then returned to 25% B at 15.1 minutes and maintained at 25% until 18 minutes. The flow rate was set to 400 μL/minute, and the column temperature was maintained at 40 °C. Mass spectrometric data were acquired over an m/z range of 220–1000 with an acquisition rate of 1 Hz.

QC samples were injected every 20 sample runs to monitor instrument performance.

#### Data processing

LC-MS data from 4-channel analysis were first exported to .csv file with Bruker DataAnalysis 4.4. The exported data were uploaded to IsoMS Pro 1.2.16. After Data Quality Check, Data Processing was performed. During feature detection and alignment, the following parameters were applied: a mass range restricted to m/z 220–1000, a saturation intensity threshold of 15,000,000, a retention time tolerance of 9 seconds, and a mass tolerance of 10 ppm.

#### Data Quality Check

Mass accuracy was checked for each sample in each of the 4 channels. Five calibration data were used to check the retention time in each channel. All calibrant peaks were well aligned. The retention times of calibrants were consistent for all calibration data, showing good retention time stability for data acquisition.

#### Data cleansing

Peak pairs without data present in at least 80.0% of samples in any experimental group were filtered out. Data were normalized by Ratio of Total Useful Signal. The missing values of peak pairs in some samples due to low signal intensity (i.e., below the detection limit) was replaced with a rationally determined ratio by a unique zero-imputation program.

#### Metabolite identification

The parameters used for metabolite identification were as follow: Retention Time Tolerance for CIL Library ID of 10 seconds, Retention Time Tolerance for LI Library ID of 75 seconds, Mass Tolerance for CIL Library ID of 10 ppm, Mass Tolerance for LI Library ID of 10 ppm, Mass Tolerance for Mass-Based Database ID of 10 ppm.

Three-tier ID approach was used to perform metabolite identification. In tier 1, peak pairs were searched against a labelled metabolite library (NovaMT Metabolite Database v3.0. CIL Library) based on accurate mass and retention time. The CIL Library contains more than 1,500 experimental entries. 280 peak pairs were positively identified in tier 1. In tier 2, linked identity library (LI Library) was used for identification of the remaining peak pairs. LI Library includes over 9,000 pathway-related metabolites, providing high-confidence putative identification results based on accurate mass and predicted retention time matches. 580 peak pairs were putatively identified in tier 2.

In tier 3, the remaining peak pairs were searched, based on accurate mass match, against the MyCompoundID (MCID) library composed of 8,021 known human endogenous metabolites (zero-reaction library), their predicted metabolic products from one metabolic reaction (375,809 compounds) (one-reaction library) and two metabolic reactions (10,583,901 compounds) (two-reaction library). 793, 1430 and 775 peak pairs were matched in the zero-, one- and two-reaction libraries, respectively. Thus, out of 4498 unique peak pairs detected, 3858 pairs could be positively identified or putatively matched. Among them, 860 peak pairs were identified as high-confidence results (tier 1 and tier 2), which were used for downstream statistical analysis.

#### Statistical analysis

Statistical analysis was performed on log-transformed data as this transformation increases the number of metabolites presenting a normal distribution. Once log-transformed, more than 90% of the metabolites were normally distributed (Shapiro-Wilk test, p < 0.01).

Principal component analysis (PCA) was performed on log-transformed data using the *pca* function in the *mixOmics* R package^82^ followed by Permutational Multivariate Analysis of Variance (PERMANOVA) using the *adonis* function in the *vegan* R package^83^. Different variables were included in the model, namely cancer, antibiotics, pair-feeding and caloric intake, to evaluate the explanatory power of each factor individually and their potential interactions.

A two-way ANOVA with Tukey’s post hoc tests on selected pairs (CT vs CT-ATB, CT vs C26, C26 vs C26-ATB) was used on log-transformed data for evaluating the ATB and C26 effects, while Student t-test was used for comparing the pair-feeding (PF) effect between PF-CT and PF-C26, when analysing bAAms (hypothesis-driven selection). When looking at the 860 peak pairs, a similar univariate approach was applied with correction for multiple testing according to the Benjamini and Hochberg (BH) procedure^84^. Only metabolites with a q-value < 0.05 for the Tukey’s post hoc test comparing C26 vs CT mice and CT-ATB vs CT mice were selected for further exploration (hypothesis-free selection).

### Chemicals

Dexamethasone and the metabolites 2-phenylpropanoic acid (2PPA, CAS 492-37-5) and pyrocatechol sulfate (PyrS, CAS 4918-96-1), were purchased from Sigma-Aldrich (USA). The other metabolites were purchased from Thermo Fisher Scientific (USA) for 3-(3,4-dihydroxyphenyl)propanoic acid (3,4OHPP, CAS 1078-61-1), from TCI Europe (Belgium) for phenylacetylglycine (PAGly, CAS 500-98-1) and synephrine (SYN, CAS 5985-28-4), and from Abcr GmBH (Germany) for N(γ)-acetyldiaminobutyric acid (NADA, CAS 1190-46-1). The PHGDH inhibitors, CBR-5884 and NCT-503, and the ‘PHGDH-inactive’ were purchased from MedChemExpress (USA).

### Cell culture

Murine colon carcinoma 26 cells (kindly provided by Dr. Mario Colombo, Fondazione IRCCS Istituto Nazionale Tumori, Italy) were maintained in DMEM high glucose medium supplemented with 10% fetal bovine serum (FBS), and 1% penicillin/streptomycin at 37°C with 5% CO_2_.

Murine C2C12 myoblasts were cultured in growth medium containing high glucose Dulbecco’s Modified Eagle Medium (DMEM) high glucose (Gibco, USA) supplemented with 7.96% Fetal Bovine Serum (Biowest, FRANCE), 0.88% MEM Non-Essential Amino Acids Solution (Gibco, USA), 88.5 U/mL penicillin-streptomycin (Gibco, USA), and 3.5 mM L-glutamine (Gibco, USA) at 37°C with 5% CO_2_. Once the C2C12 cells reached 80-90% confluence, growth medium was replaced with differentiation medium containing DMEM high glucose (Gibco, USA) supplemented with 1.88% Horse Serum (Gibco, USA), 0.94% MEM Non-Essential Amino Acids Solution (Gibco, USA), 100 U/mL penicillin-streptomycin (Gibco, USA), and 4 mM L-glutamine (Gibco, USA). After four days of differentiation, cells were treated for 48 h with vehicle or dexamethasone at 1 µM, with or without the metabolites (individually or in combination) at 10 or 100 µM, with or without PHGDG inhibitor (CBR-5884 ranging from 2.5 to 33 µM, or NCT at 2.5 or 10 µM), with or without the PHGDH-inactive control (2.5 or 10 µM). For myotube diameter quantification, images were captured using a phase contrast microscopy (EVOS XL Core or EVOS M3000 Imaging System, Thermo Fisher Scientific, Belgium). The myotube diameter was quantified blindly with the image processing software ImageJ (U.S. National Institutes of Health, USA). Four pictures were taken of each culture dish and the experiments were performed in technical triplicates and, when indicated in the legends, also in biological triplicates. For each picture, 10 myotubes were randomly selected and five measurements for each myotube were performed. Myotubes were defined by the presence of a minimum of 5 nuclei.

### LDH assays

To assess the cytotoxicity of the treatment on the myotubes, the lactate dehydrogenase released in the supernatant was measured using a biochemical assay (LDH 21 FS, DiaSys Diagnostic Systems, Germany) following the manufacturer’s instructions. Cytotoxicity was expressed relative to the positive control, Triton X-100 (1%), which was defined as inducing 100% cytotoxicity. Sample responses were normalized accordingly. A condition causing a cytotoxicity value exceeding a threshold of 10% relative to the positive control was considered cytotoxic.

### Transcriptome analysis

Total RNA was isolated from C2C12 myotubes by TriPure reagent (Roche, Switzerland). Samples were treated with DNase using the DNA-free™ kit (Ambion, USA), and RNA quantity was evaluated using a NanoPhotometer Spectrophotometer (Implen, Germany). The quality of the RNA samples was assessed using a 2100 Bioanalyzer System (Agilent Technologies, USA). All RIN values were above 7, supporting RNA integrity. Illumina TruSeq Stranded mRNA libraries were prepared using polyA tail selection. Following library quality control, libraries were pooled and sequenced using a 2 × 150bp paired-end configuration on a NovaSeq X instrument (Macrogen, The Netherlands). Two samples (one CT and one Combi_10) were excluded due to the low quality of their library. FASTQ files were processed using a standard RNAseq pipeline including Trimmomatic (version 0.39)^85^ to remove low quality reads and HISAT2 (version 2.2.1)^86^ to align reads to the mouse reference genome (GRCm38). Gene expression levels were quantified using featureCounts from Subread (version 2.0.3)^87^ and genomic features were defined based on the Ensembl Mus_musculus.GRCm38.94.gtf annotation file. PCA was performed on log-transformed data using the *prcomp* function on rlog-transformed gene expression data, based on the 500 genes with the highest variance across samples. Differential expression analyses were performed with DESeq2 Bioconductor package (version 1.36.0)^88^. Genes with an adjusted p-value < 0.05 were considered differentially expressed. Over-representation analyses (ORA) were performed using clusterProfiler (version 4.14.4)^89^ on genesets from the molecular Signature Database (MSigDB) downloaded using msigdbr package (version 7.5.1). Prediction of transcription factors involved in gene network regulation was carried out using human databases from TRRUST, as implemented in Metascape (version 3.5) with a separate analysis for genes upregulated and genes downregulated^90^.

### Statistical analysis

When evaluating the impact of each individual metabolite *in vitro*, paired Student t-test was performed. For subsequent analyses, a one-way ANOVA was performed followed by Dunnett’s pairwise comparison post-test to the reference group (either vehicle or dexamethasone alone, as appropriate). In the case where selected pairs of post-test comparisons were performed, one-way ANOVA was followed by Sidak’s multiple comparisons test. When comparing more than two groups and assessing two independent factors (2PPA and 3,4OHPP effect), a two-way ANOVA was performed. Metabolomics data analyses are described in the Metabolomics section. In any case, p < 0.05 was considered statistically significant. Data are presented as mean ± SEM. Statistical analyses were carried out using GraphPad Prism v8.0.1 for windows (GraphPad Software, USA) and R.

## DATA AVAILABILITY STATEMENT

Raw transcriptomic data can be found on Gene Expression Omnibus (GEO) public repository (GSE343204, will be made public upon publication). The remaining data supporting the findings of this study are available within the article and its supplementary materials.

## ACKNOWLEDGMENTS

We are grateful to Stéphanie Delieux and Bouazza Es Saadi for their precious and skilled technical assistance throughout the entire course of this work. We also thank Dr Adeline Dolly, Sophie Lecop and Savannah Eeckhout for their help with preliminary experiments, mouse husbandry and necropsy, as well as Dr Sarah Pötgens for fruitful discussion around the experimental design of the *in vivo* experiment and associated metabolomics. We are grateful to Prof Jean-Paul Thissen, Dr Isabelle Massart and Pascal Lause for guidance with C2C12 myotubes culture and analyses. We thank Prof Liang Li (University of Alberta, Canada, The Metabolomics Innovation Centre) and his team for generating metabolomics data. We benefited from the access to and expertise of the Ghent Germ-free and Gnotobiotic mouse facility, for which we thank Prof Lars Vereecke and Dr Vanessa Andries.

## AUTHOR CONTRIBUTION

Conception and design of the work: LBB. Experimental designs: ALD, XL, AL, LBB. Sample and data collection: ALD, XL, CL. Setup of the dexamethasone model and assistance with *in vitro* experiments: EP. Transcriptomics analyses: AL. Metabolomics analysis: LBB. Other data processing: ALD, XL. Statistical analyses: ALD, AL, LBB. Data interpretation: ALD, XL, LBB. Participation to critical scientific discussions: NMD, AMN. Acquisition of funding: LBB. Drafting the article: ALD, LBB, with contribution from XL and AMN for the draft of the figures. Critical revision of the article: all.

