## Supplementary Figures File for "Gut microbiota-dependent phenylpropanoic acid derivatives reduced in cancer cachexia protect against myotube atrophy"

### Supplemental Figure 1

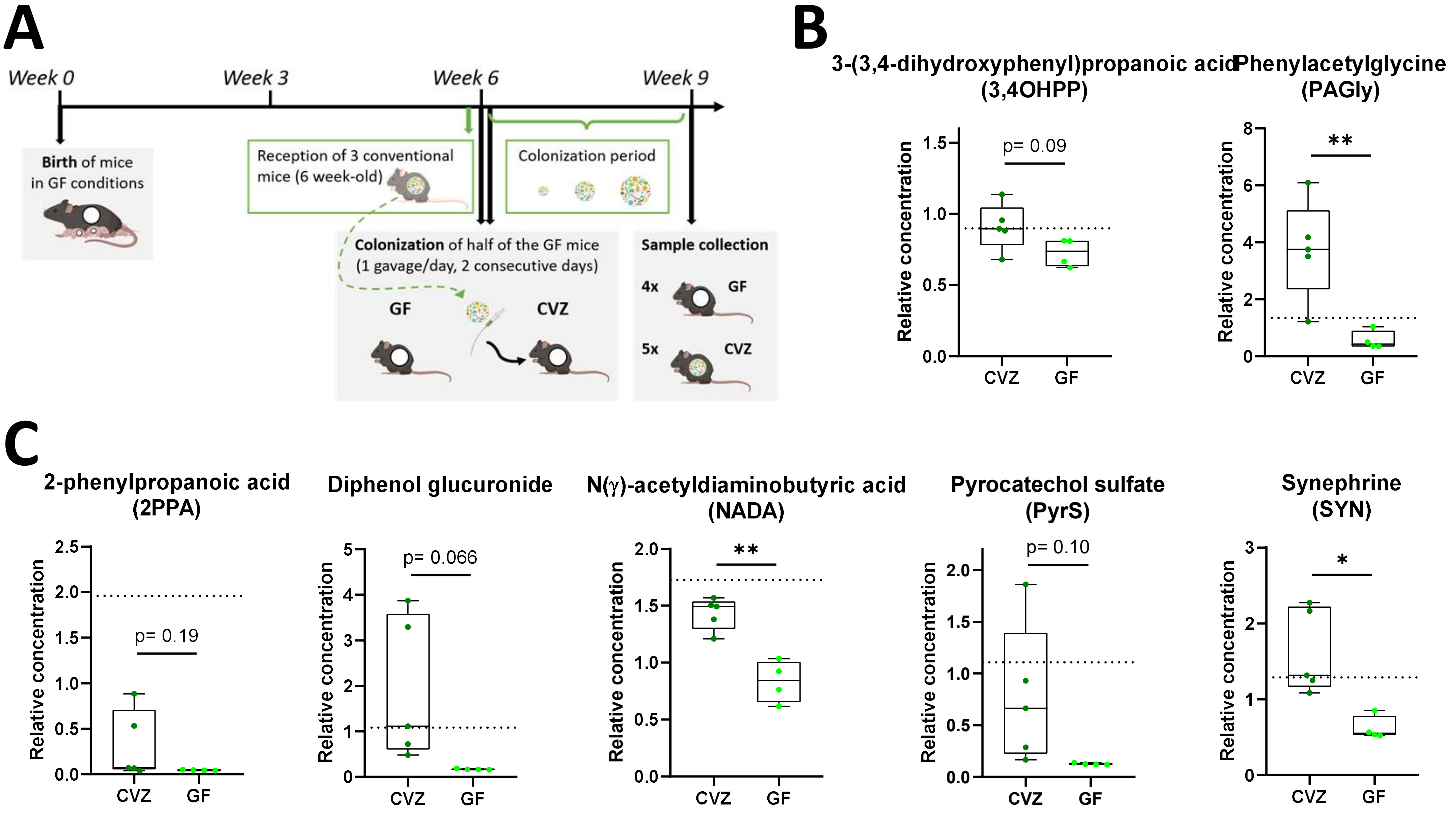


**Figure S1. Selected metabolites are increased upon conventionalization of germ-free mice.** (**A**) Schematic overview of the experimental design. Mice born germ-free (GF) were divided in two groups: mice maintained in GF conditions and GF mice conventionalized by gavage with the microbiota of conventional mice (CVZ). Blood samples from CVZ and GF mice were subjected to metabolite screening by chemical isotope labelling liquid chromatography–mass spectrometry (CIL LC–MS). (**B**) Relative concentration of metabolites selected based on a hypothesis-driven approach: 3-(3,4-dihydroxyphenyl)propanoic acid (3,4OHPP) and phenylacetylglycine (PAGly). (**C**) Relative concentration of metabolites selected using a hypothesis-free approach: 2-phenylpropanoic acid (2PPA), diphenol glucuronide, N(γ)-acetyldiaminobutyric acid (NADA), pyrocatechol sulfate (PyrS), and synephrine (SYN). Data represented in **B** and **C** were analysed by Student t-test to compare CVZ vs GF mice. Dotted horizontal line indicates the mean value obtained in the CT mice presented in Figure 1. N = 4-5 mice per group. *$p \leq 0.05$ and **$p \leq 0.01$.

### Supplemental Figure 2

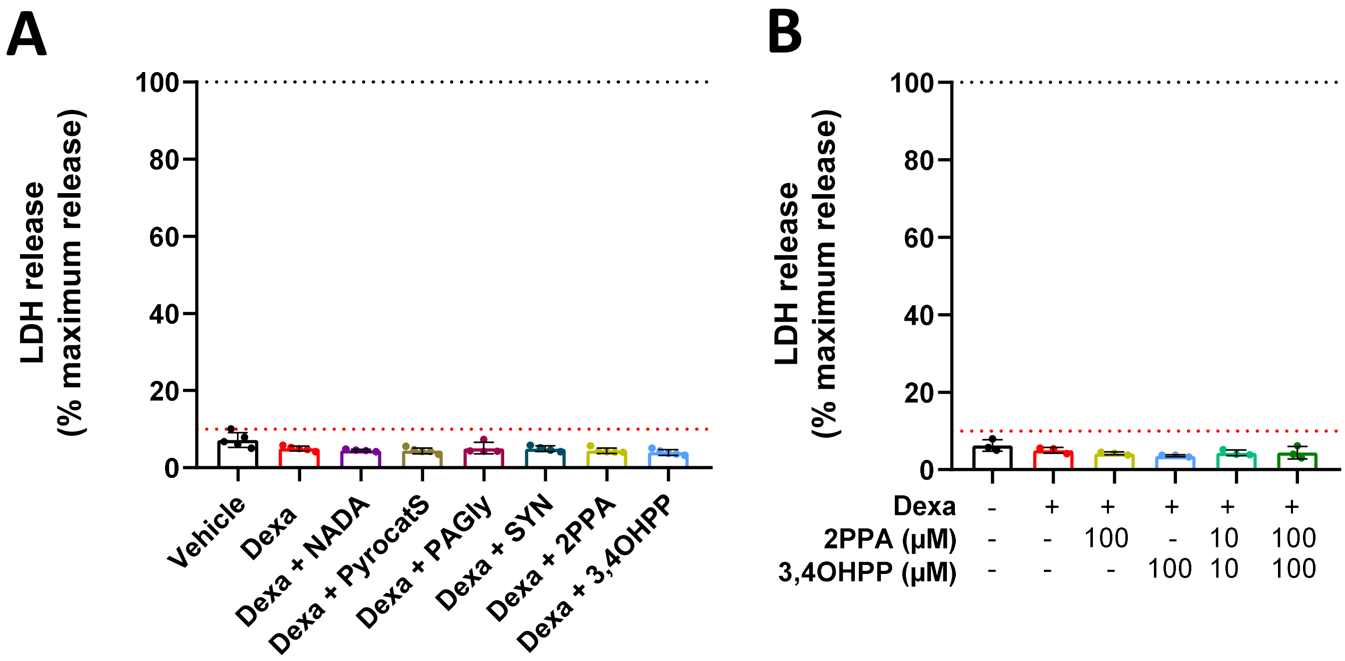


**Figure S2. Selected bacteria-dependent metabolites attenuate dexamethasone-induced muscle atrophy in C2C12 myotubes.** C2C12 myotubes were treated with different conditions for 48 h and cytotoxicity level was measured. (**A**) In the presence of each individual candidate metabolites (100 µM) with the presence of 1 µM dexamethasone (Dexa). N = 5-8 experiments, n = 3 technical replicates. (**B**) Following combined treatment with 2PPA and 3,4OHPP at 10 or 100 µM, in the presence of 1 µM Dexa. N = 4 experiments, n = 3 technical replicates. Data were analysed relatively to cells treated with Triton (positive control, black dotted line). Red dotted line represents the cytotoxicity threshold.

### Supplemental Figure 3

A

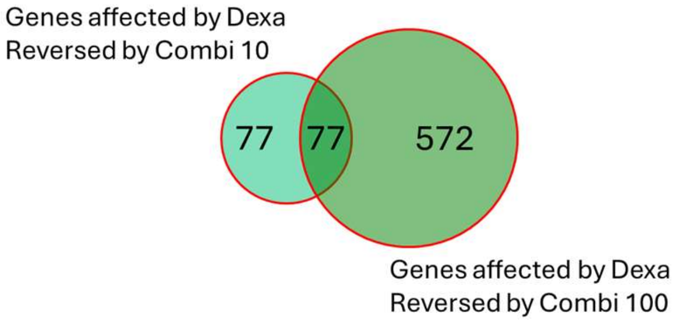


B

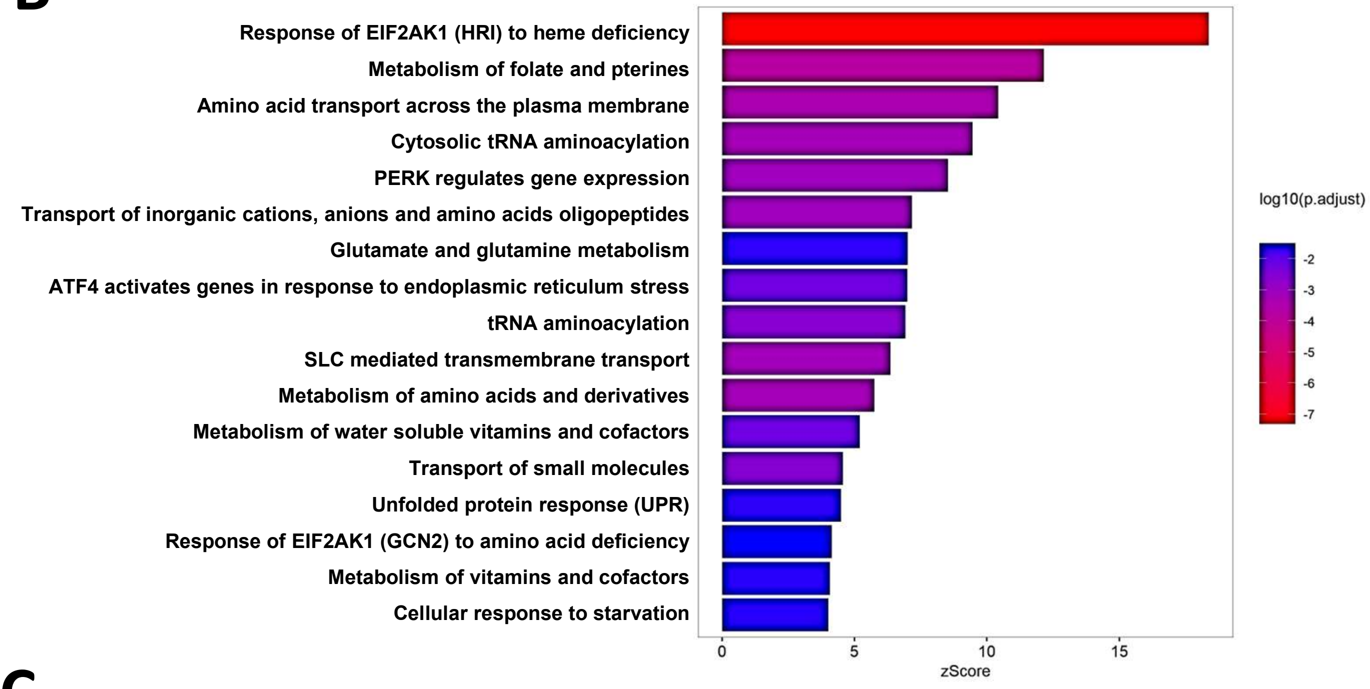


C

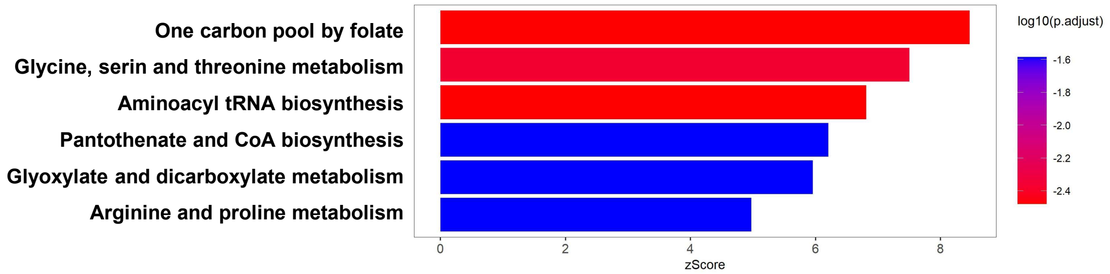


D

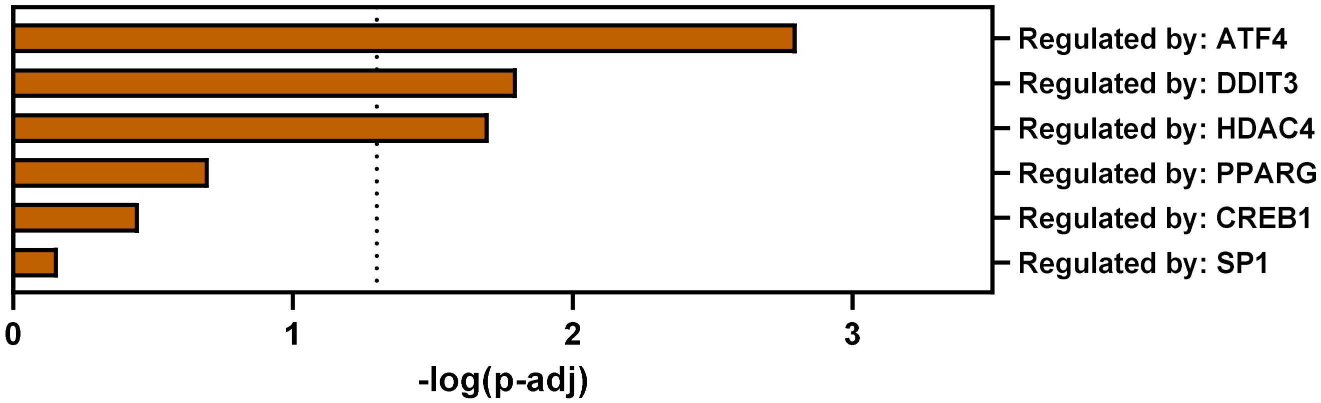

**Figure S3. Genes altered by dexamethasone and reversed by the combination of 2PPA and 3,4OHPP at 10µM and 100µM are involved in amino acid homeostasis and under the control of ATF4.** (**A**) Venn diagram showing the number of genes significantly affected by dexamethasone and whose changes are reversed upon addition of a combination of 2PPA and 3,4OHPP at 10 µM (left) or 100 µM (right). 77 genes are reversed at both concentrations. (**B-C**) Significantly enriched REACTOME (**B**) and KEGG (**C**) pathways identified through over-representation analysis (ORA) of these 77 genes. (**D**) Transcription factors predicted to be involved in the regulation of these 77 genes using TTRUST implemented in Metascape. All transcription factors with a p-value < 0.05 are depicted. The vertical dotted line indicates the significant threshold (p-adj ≤ 0.05). N = 1 experiment, n= 6 technical replicates.

### Supplemental Figure 4

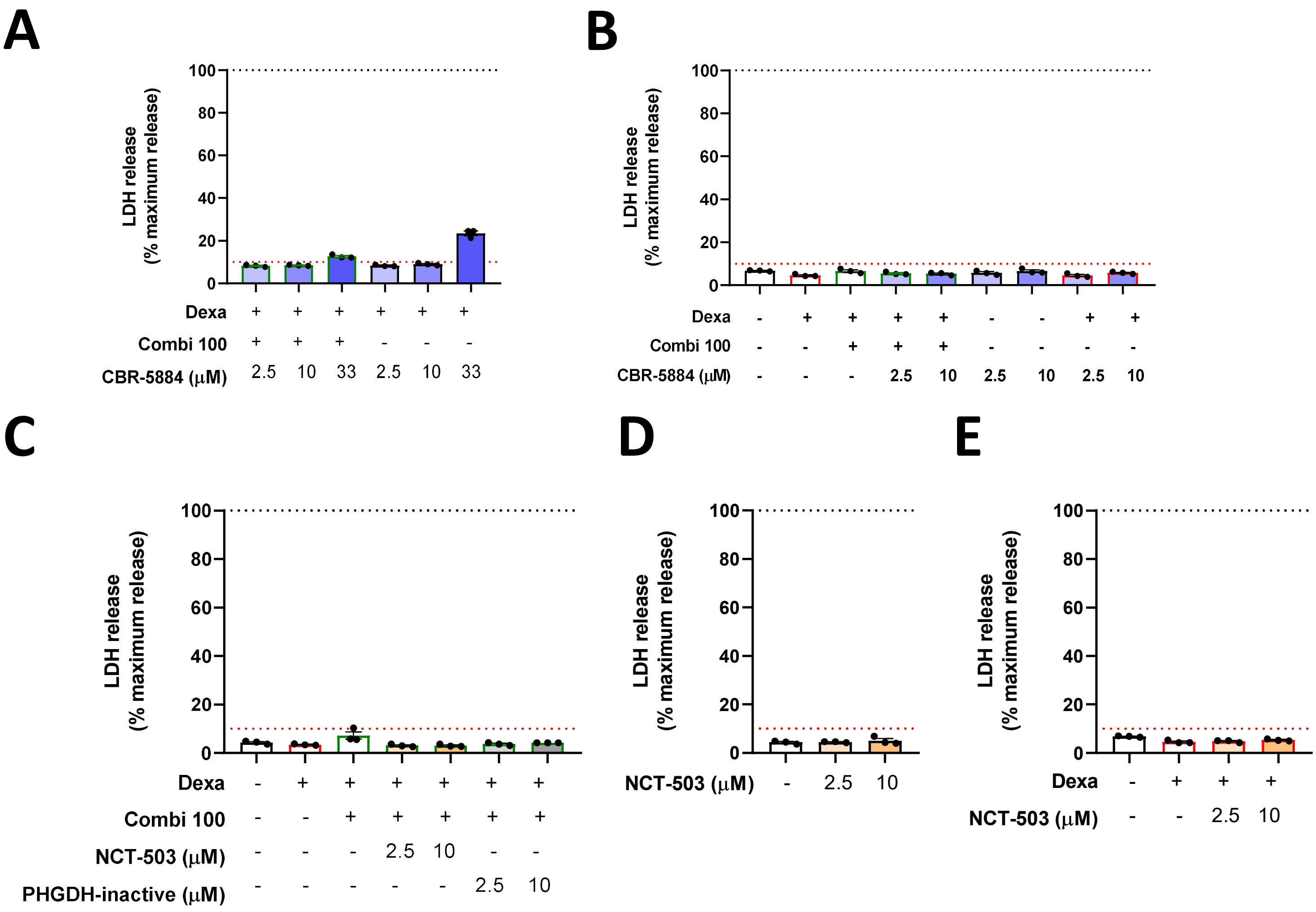


**Figure S4. Inhibition of PHGDH attenuates the protective effect of combined metabolites against dexamethasone-induced myotube atrophy.** C2C12 myotubes were treated with different conditions for 48 h and cytotoxicity level was measured. (**A**) In the presence of 1 µM dexamethasone (Dexa) and the PHGDH inhibitor CBR-5884 (2.5, 10, 33 µM) with or without combination of 2PPA and 3,4OHPP (Combi 100 µM). N = 1 experiment, n = 3 technical replicates. (**B**) In the presence or absence of 1 µM Dexa, combined or not with Combi 100 µM, and with or without the PHGDH inhibitor CBR-5884 (2.5, 10 µM). N = 3 experiments, n = 3 technical replicates. (**C**) In the presence or absence of 1 µM Dexa, combined or not with Combi 100 µM, with or without the PHGDH inhibitor NCT-503 (2.5, 10 µM), and with or without PHGDH-inactive (2.5, 10 µM). N = 3 experiments, n = 3 technical replicates. (**D**) In the presence or absence of the PHGDH inhibitor NCT-503 (2.5, 10 µM). N = 3 experiments, n = 3 technical replicates. (**E**) In the presence or absence of 1 µM Dexa, combined or not the PHGDH inhibitor NCT-503 (2.5, 10 µM). N = 3 experiments, n = 3 technical replicates. Data were analysed relatively to cells treated with Triton (positive control, black dotted line). Red dotted line represents the cytotoxicity threshold.
