## Supplementary Table File for "Gut microbiota-dependent phenylpropanoic acid derivatives reduced in cancer cachexia protect against myotube atrophy"

### Supplementary Table 1. List of screened metabolites for the hypothesis-driven approach.

Bacterial amino acid metabolites (bAAms) were identified through a literature review (Lefevre & Bindels, *Curr Osteoporos Rep* 2022 ; Oliphant & Allen-Vercoe, *Microbiome* 2019 ; Nemet et al, *Cell* 2020), and chemical analogs were selected too.

| **Metabolite list, selected based on a litterature search focussing on bacterial metabolites from amino acids** | **Metabolite list, selected in the metabolomics dataset based on their structural homology with known bAAms, based on the "hydroxyphenyl" name** |
|---|---|
| 2,4-diaminopentanoic acid | 2-hydroxy-3-(4-hydroxyphenyl)propenoic acid |
| 2-methylbutyrate/2-methylbutanoic acid | 2-hydroxyphenylacetic acid |
| 2-methylbutyrylglycine | 2-hydroxyphenylacetic acid O-sulfate |
| 3-phenylpropionic acid | 3-(2,3-dihydroxyphenyl)propanoic acid |
| 4-aminobutyrate (gamma-aminobutyrate) | 3-(3,4-dihydroxyphenyl)lactic acid |
| 4-hydroxyphenyl-acetaldehyde | 3-(3,4-dihydroxyphenyl)pyruvic acid |
| 4-hydroxyphenyl-acetate | 3-(3-hydroxyphenyl)propionic acid |
| 4-hydroxyphenyl-acetate-taurine | 3,4-dihydroxyphenylacetaldehyde |
| 4-hydroxyphenyl-acetylglycine/p-hydroxyphenylacetylglycine | 3-(3,4-dihydroxyphenyl)propanoic acid |
| 4-hydroxyphenyl-butyrate | 3-amino-3-(4-hydroxyphenyl)propanoic acid |
| 4-hydroxyphenyl-butyrate-taurine | 3-hydroxyphenylacetic acid |
| 4-hydroxyphenyl-butyrylglycine | 3-hydroxyphenylpyruvic acid |
| 4-hydroxyphenyllactate | |
| 4-hydroxyphenyl-lactate-glycine | |
| 4-hydroxyphenyl-lactate-taurine | |
| 4-hydroxyphenyl-propionate | |
| 4-hydroxyphenyl-propionate-taurine | |
| 4-hydroxyphenyl-propionylglycine | |
| 4-hydroxyphenylpyruvate | |
| 4-hydroxyphenyl-pyruvate-glycine | |
| 4-hydroxyphenyl-pyruvate-taurine | |
| 5-acetamidopentanoic acid | |
| 5-aminopentanal | |
| 5-aminopentanamide | |
| 5-aminopentanoic acid | |
| acetate | |
| agmatine (4-guanidinobutanamide) | |
| butyrate | |
| cadaverine | |
| cresyl-sulfate (p-cresol sulfate) | |
| gamma-L-glutamylputrescine | |
| histamine | |
| imidazole acetic acid | |
| imidazole lactic acid | |
| imidazole propionate | |
| imidazole pyruvic acid | |
| indole | |
| indole-3-acetaldehyde | |
| indole-3-acetamide | |
| indole-3-acetic acid (indoleacetate) | |
| indole-3-aldehyde (3-formylindole) | |
| indole-3-lactic acid (indolelactate) | |
| indole-3-sulfate (indoxyl sulfate) | |
| indoleacrylic acid | |
| isobutyrate | |
| isobutyrylglycine | |
| isovalerate | |
| methylamine | |
| N-acetylindoxyl | |
| p-coumarate (4-hydroxycinnamate) | |
| p-coumaroylputrescine | |
| p-cresol (4-cresol) | |

| |
|---|
| phenol<br>phenylacetic acid<br>phenylacetylglycine<br>phenylacetyltaurine<br>phenylbutyrate<br>phenylbutyrylglycine<br>phenylethylamine<br>phenyllactate<br>phenyl-lactate-glycine<br>phenyl-lactate-taurine<br>phenylpropionylglycine<br>phenylpropionyltaurine<br>phenylpyruvate<br>phenyl-pyruvate-glycine<br>phenyl-pyruvate-taurine<br>propionate<br>putrescine (1,4-diaminobutane)<br>spermidine<br>spermine<br>succinate<br>trans-2,3-dihydroxycinnamic acid<br>trans-cinnamic acid<br>tryptamine<br>tryptophol (indole-3-ethanol)<br>tyramine<br>urocanate |
